# Norepinephrine and dopamine contribute to distinct repetitive behaviors induced by novel odorant stress in male and female mice

**DOI:** 10.1101/2022.01.25.477573

**Authors:** Daniel J. Lustberg, Joyce Q. Liu, Alexa F. Iannitelli, Samantha O. Vanderhoof, L. Cameron Liles, Katharine E. McCann, David Weinshenker

**Author notes:** **Conflict of interest statement**: DW is co-inventor on a patent concerning the use of selective dopamine β-hydroxylase inhibitors for the treatment of cocaine dependence (US-2010-0105748-A1; “Methods and Compositions for Treatment of Drug Addiction”). The other authors declare no conflicts of interest. **Author approvals**: all authors have seen and approved the manuscript, and it hasn’t been accepted or published elsewhere.

## Abstract

Exposure to unfamiliar odorants induces an array of repetitive defensive and non-defensive behaviors in rodents which likely reflect adaptive stress responses to the uncertain valence of novel stimuli. Mice genetically deficient for dopamine β-hydroxylase (*Dbh−/−*) lack the enzyme required to convert dopamine (DA) into norepinephrine (NE), resulting in globally undetectable NE and supranormal DA levels. Because catecholamines modulate novelty detection and reactivity, we investigated the effects of novel plant-derived odorants on repetitive behaviors in *Dbh−/−* mice and *Dbh+/−* littermate controls, which have catecholamine levels comparable to wild-type mice. Unlike *Dbh+/−* controls, which exhibited vigorous digging in response to novel odorants, *Dbh−/−* mice displayed excessive grooming. Drugs that block NE synthesis or neurotransmission suppressed odorant-induced digging in *Dbh+/−* mice, while a DA receptor antagonist attenuated grooming in *Dbh−/−* mice. The testing paradigm elicited high circulating levels of corticosterone regardless of *Dbh* genotype, indicating that NE is dispensable for this systemic stress response. Odorant exposure increased NE and DA abundance in the prefrontal cortex (PFC) of *Dbh+/−* mice, while *Dbh−/−* animals lacked NE and had elevated PFC DA levels that were unaffected by novel smells. Together, these findings suggest that novel odorant-induced increases in central NE tone contribute to repetitive digging and reflect psychological stress, while central DA signaling contributes to repetitive grooming. Further, we have established a simple method for repeated assessment of stress-induced repetitive behaviors in mice, which may be relevant for modeling neuropsychiatric disorders like Tourette syndrome or obsessive-compulsive disorder that are characterized by stress-induced exacerbation of compulsive symptoms.

## 1. Introduction

In rodents, exposure to novel or noxious stimuli elicits increases in repetitive and defensive behaviors as well as circulating levels of the stress hormone corticosterone (CORT) (Bowers et al., 2008; Garbe et al., 1993; Kemble and Bolwahnn, 1997; Seggie and Brown, 1975). Various psychological stressors also facilitate the release of the catecholamine neuromodulators norepinephrine (NE) and dopamine (DA) in regions like the prefrontal cortex (PFC) (Abercrombie et al., 1989; Aston-Jones et al., 1996; Finlay et al., 1995; Valentino et al., 1993), which is implicated in stress-induced anxiety and the development of repetitive and stereotyped behaviors (Ahmari and Rauch, 2022; Kenwood et al., 2022; McKlveen et al., 2015; Shansky and Lipps, 2013). Neurons in the PFC receive both NE inputs from the brainstem locus coeruleus (LC) and DA inputs from the midbrain ventral tegmental area (VTA) (Pan et al., 2004; Shansky and Lipps, 2013), and the firing rate of NE and DA projections to the PFC are altered by various stressors (Aston-Jones et al., 1996; Morilak et al., 2005; Valenti et al., 2011). Thus, PFC catecholamine tone could represent a sensitive index of psychological stress, and fluctuations in PFC NE and DA signaling may drive stress-induced anxiety-like and repetitive behaviors (Dzirasa et al., 2010; Iimori et al., 1982; Lustberg et al., 2020a; Lustberg et al., 2020b; McKlveen et al., 2015; Shah et al., 2004).

Plant-derived odorants from essential oils are novel olfactory stimuli for laboratory mice that we posited would engender similar behavioral manifestations of psychological stress as spatial novelty or innately threatening olfactory stimuli like predator-derived scents (Dielenberg and McGregor, 2001; Hebb et al., 2002; Hwa et al., 2019; Kemble and Bolwahnn, 1997; Kemble et al., 1995; Lustberg et al., 2020b). Because many plants contain toxic secondary metabolites and/or have little or no nutritional value for rodents, defensive and avoidant responses to unfamiliar plant-derived olfactory stimuli would be adaptive for rodents in the wild to discourage appetitive behavior (Garbe et al., 1993; Hansen et al., 2016; Kemble and Gordon, 1996). Indeed, novel non-predator odorants elicit robust activation of risk assessment behaviors in mice (e.g. defensive burying, stretch attend posture, flat-back approach, etc.) and suppress consummatory and passive behaviors (e.g. eating, grooming) (Garbe et al., 1993; Kemble and Bolwahnn, 1997; Kemble and Gibson, 1992). Defensive behaviors, including burying in the shock probe test and digging in the marble burying (MB) assay, are facilitated by central NE neurotransmission (Bondi et al., 2007; De Boer and Koolhaas, 2003; den Hartog et al., 2020; Howard et al., 2008; Lustberg et al., 2020a), while many consummatory and non-defensive behaviors such as grooming are mediated by central DA transmission (Blackburn et al., 1992; Kiyatkin, 1995; Murray and Waddington, 1989). Thus, one goal of these experiments was to further parse the specific contributions of NE and DA to repetitive behavioral responses elicited by novel odorants.

Dopamine β-hydroxylase (DBH) converts DA to NE, thus controlling catecholamine ratios and content in noradrenergic neurons and their target regions. We have previously shown that global DBH knockout (*Dbh−/−*) mice, which completely lack NE and have elevated levels of central and peripheral DA (Bourdélat-Parks et al., 2005; Thomas et al., 1998; Thomas et al., 1995), are behaviorally insensitive to stress typically induced by novel environments or cage change in multiple assays, including novelty-suppressed feeding, nestlet shredding, and MB tests (Lustberg et al., 2020a; Lustberg et al., 2020b). These anxiety-like and repetitive behaviors were rescued to control levels in *Dbh−/−* mice by transient pharmacological restoration of central NE synthetic capacity, and the *Dbh*−/− behavioral phenotypes were mimicked in NE-competent *Dbh+/−* control mice using systemic administration of drugs that reduce central NE synthesis, release, or adrenergic receptor (AR) signaling (Lustberg et al., 2020a; Lustberg et al., 2020b). The absence of species-typical, innate behavioral responses to stress elicited by spatial novelty or cage change in *Dbh−/−* mice was associated with hypoactivity of forebrain targets of the LC including the PFC, which is highly implicated in the expression of compulsive behavior and stress-induced anxiety (Ahmari and Rauch, 2022; Arnsten et al., 2017; Koga et al., 2020; McKlveen et al., 2015; Shah and Treit, 2003).

Given the behavioral indifference of *Dbh−/−* mice to novel environments and tastes (Lustberg et al., 2020b), we hypothesized that exposure to novel odorants would produce a dissociation in stress-related defensive responses between *Dbh−/−* and *Dbh+/−* mice. Novelty is a quality of stimuli in all sensory modalities; behavioral responses to novelty in rodents can be further subdivided into novelty detection, which engages memory circuits, and novelty reactivity, which engages emotional circuits (Dellu et al., 1996; Fanselow and Dong, 2010; Kafkas and Montaldi, 2018; Lee et al., 2005; Stone et al., 1999). In the current study, we assessed the expression of repetitive defensive (digging) and non-defensive (grooming) behaviors in *Dbh−/−* mice and their NE-competent *Dbh+/−* littermate controls in the presence of unfamiliar plant-derived odorants. Next, we assessed the behavioral effects of drugs that interfere with NE synthesis, release, or signaling in *Dbh+/−* mice and DA receptor antagonism in *Dbh−/−* mice exposed to the novel plant-derived odorants. Finally, we assessed central catecholamine tissue content and tyrosine hydroxylase (TH) phosphorylation status in the PFC and the dorsal pons/LC as well as circulating levels of the stress hormone CORT following novel odorant exposure to determine which, if any, of these physiological indices were associated with reactivity to olfactory novelty stress (Dunkley and Dickson, 2019; McKlveen et al., 2015).

## 2. Materials and methods

### 2.1 Mice and housing

*Dbh*−/− mice were maintained on a mixed 129/SvEv and C57BL/6J background, as previously described (Thomas et al., 1998; Thomas et al., 1995). Pregnant *Dbh*+/− dams were administered drinking water containing the β-adrenergic receptor (AR) agonist isoproterenol and the α1AR agonist phenylephrine (20 μg/ml each; Sigma-Aldrich) with vitamin C (2 mg/ml) from E9.5–E14.5, and the synthetic NE precursor L-3,4-dihydroxyphenylserine (DOPS; 2 mg/ml; Lundbeck, Deerfield, IL) + vitamin C (2 mg/ml) from E14.5-parturition to prevent embryonic lethality resulting from complete *Dbh* deficiency (Mitchell et al., 2008; Thomas et al., 1995). *Dbh*−/− mice are easily distinguished from their NE-competent littermates by their visible delayed growth and bilateral ptosis phenotypes, and genotypes were subsequently confirmed by PCR. *Dbh*+/− littermates were used as controls because their behavior and catecholamine levels are indistinguishable from wild-type (*Dbh+/+*) mice (Bourdélat-Parks et al., 2005; Szot et al., 1999; Thomas et al., 1998).

A total of 51 male and female mice (8–16 months of age) were used for all experiments. The groups consisted of 28 *Dbh*+/− (n = 22 male, n = 12 female) and 17 *Dbh*−/− (n = 7 male, n = 10 female) animals. No sex differences in stress-induced digging or grooming were observed in pilot experiments and have not been reported in the literature (Dixit et al., 2020; Li et al., 2006; Londei et al., 1998; Smolinsky et al., 2009); thus, male and female mice of the same *Dbh* genotype were pooled and evenly distributed across groups. All animal procedures and protocols were approved by the Emory University Animal Care and Use Committee in accordance with the National Institutes of Health guidelines for the care and use of laboratory animals. Mice were maintained on a 12 h/12 h light/dark cycle (7:00/19:00 h) with access to food and water *ad libitum* except during behavioral testing. Behavioral testing was conducted under standard lighting conditions during the light cycle (ZT5-ZT8) in the same room where the mice were housed to minimize the stress of cage transport on test days.

### 2.2. Behavioral pharmacology experiments with novel odorants and odorless control

Mice were group-housed in static cages in same-sex groups of 2-5. On test days, individual mice were removed from their home cages and placed into new standard mouse cages (13” x 7” x 6”) containing only clean bedding substrate and a cotton nestlet square (5 cm x 5 cm, ~3g) pre-treated with 1 ml of either deionized water (odorless control) or a novel plant odorant dissolved 1:100 in deionized water. The plant-based essential oils employed in this study came from a kit (Anjou, Fremont, CA) that included lavender, tea tree, patchouli, cypress, and palmarosa. A clear plexiglass cover was pressed flush over the cages to prevent vaporization or dispersal of the odorants over the course of testing while still allowing filming and behavioral scoring. The mice in the odorant-treated cages were filmed with a front-facing, mounted digital video camera for 10 min for assessment of repetitive behaviors including defensive (digging/burying) and non-defensive (grooming) behaviors (De Boer and Koolhaas, 2003; Kalueff et al., 2016; Kemble and Bolwahnn, 1997; Londei et al., 1998; Pond et al., 2021; Smolinsky et al., 2009). Mice were returned to their home cages after the 10-min odorant exposure and were never re-exposed to the same odorant twice. Cage mates were used in “batches” to prevent premature odorant exposure and possible olfactory cross-contamination. Time spent digging and grooming (sec) during the 10-min task were manually scored by trained observers using digital stopwatches.

### 2.2.1. Assessing behavioral responses to cage change with odorless stimulus or novel odorant

*Dbh−/−* (n = 7) and control (n = 6) mice of both sexes (~8 months old) were assessed for defensive and non-defensive repetitive behavioral responses following exposure to water or a novel plant-derived odorant in a new clean cage. Mice of both genotypes were first tested in the cage change + water condition, and behavior was scored for 10 min. On the following day, mice were tested in the cage change + novel odorant condition (tea tree oil) and behavior was filmed for 10 min and scored. Digging behavior was further subdivided into exploratory or “light” digging and defensive burying behaviors for this experiment only; composite digging scores are reported for pharmacological experiments in which all mice were exposed to a clean cage scented with a novel odorant.

### 2.2.2 Pharmacological manipulation of repetitive behaviors induced by novel odorants

To determine which ARs are required for the expression of repetitive digging following exposure to a clean cage scented with a novel odorant, control mice were administered i.p. injections of bacteriostatic saline (vehicle control; n = 21), the inhibitory α_2_-autoreceptor agonist guanfacine (0.3 mg/kg; n = 10) (Cayman Chemical, Ann Arbor, MI), a cocktail of the β_1/2_AR antagonist propranolol (5 mg/kg) (Sigma-Aldrich, St. Louis, MO) and the α_1_AR antagonist prazosin (0.5 mg/kg; n = 11) (Sigma-Aldrich), or the DBH inhibitor nepicastat (100 mg/kg; n = 5) (Synosia Therapeutics, Basel, Switzerland). Mice were never exposed to the same drug or odorant more than once. *Dbh−/−* mice were treated with either saline or the non-selective DA receptor antagonist flupenthixol (0.25 mg/kg) (Sigma-Aldrich) to determine if excessive DA signaling contributes to grooming in these mice.

All drugs were dissolved in sterile saline (0.9% NaCl) except for prazosin and propranolol, which were first dissolved in DMSO and Cremophor EL prior to addition to saline (final concentrations 1.5% DMSO and 1.5% Cremophor EL). All compounds were administered by i.p. injections at a volume of 10 ml/kg 30 min prior to testing except nepicastat, which was administered 2 h prior, as previously described (Lustberg et al., 2020a; Schroeder et al., 2013). Sterile saline vehicle was injected to control for the effect of injection stress on behavior, and vehicle-treated animals were used for statistical comparison with anti-adrenergic compounds. Doses selected were based on previous studies and pilot experiments to control for confounding effects such as sedation (Lustberg et al., 2020a; Mitchell et al., 2008).

### 2.3 Novel odorant exposure for physiological measurements of stress

Individual mice of both genotypes were either transferred to a new clean cage without a novel odorant present to control for the known stress effects of cage change (n = 7 *Dbh+/−* mice, n = 7 *Dbh−/−* mice) or were transferred to a new cage containing a nestlet treated with a novel plant odorant (palmarosa oil; n = 7 *Dbh+/−* mice and 9 *Dbh−/−* mice) (Balcombe et al., 2004; Rasmussen et al., 2011). The exposure lasted 10 min, after which the nestlet was removed and the cage was briefly allowed to air out. This process was also performed in the odorant-free, cage change only condition. Twenty min after the odorant exposure test ended, mice were transferred to a nearby procedure room for tissue and plasma collection. A separate experiment was conducted using only *Dbh+/−* mice treated 2 h prior with saline vehicle (n=7) or nepicastat (100 mg/kg; n = 7) prior to cage change + novel odorant (palmarosa oil) to quantify the extent of catecholamine disruption and determine if acute DBH inhibition altered other physiological measures of stress induced by novel odorant exposure.

#### 2.3.1. Plasma collection and corticosterone measurement

Mice used for corticosterone (CORT) measurement were exposed for 10 min to either an odorless clean cage or clean cage in the presence of a novel odorant and were euthanized 20 min later (30 min from onset of cage change with or without odorant exposure). Plasma was collected during the light cycle in a 2 h time window beginning 5 h after the onset of the light cycle and ending within 5 h of the onset of the dark cycle (ZT5-ZT7) to control for circadian variation in CORT. All saline- and nepicastat-treated *Dbh+/−* mice were exposed to the cage change with novel odorant condition but were euthanized in an identical fashion. Mice were anesthetized with isoflurane (Patterson Veterinary Supply, Devens, MA) and rapidly decapitated. Trunk blood was collected in EDTA-coated tubes (Sarstedt Inc., Newton, NC) and chilled on ice, as previously described (Tillage et al., 2021). Blood was centrifuged for 20 min at 3000 rpm at 4°C to isolate plasma, which was collected and stored at −80°C. CORT was measured using the Corticosterone Competitive ELISA kit (Thermo Fisher, #EIACORT, Waltham, MA) following manufacturer’s instructions for small volume blood plasma. The plate was read at 405 nm with correction at 595 nm. The cross reactivity for the assay was 12.3% for deoxycorticosterone and <0.8% for all other hormones. The inter-assay variability was 7.9% for all plates and the intra-assay variability was 5.2%. The sensitivity of the assay was 18.6 pg/ml.

#### 2.3.2. Pons and PFC dissection and HPLC detection of catecholamines and metabolites

Mice were euthanized for trunk blood and brain tissue collection following exposure to either cage change or cage change + novel odorant, as previously described and delineated above (Acosta et al., 2022; Tillage et al., 2020a). Dorsal pons (containing LC) and PFC sections were rapidly dissected on ice and snap-frozen in isopentane on dry ice. The samples were weighed and stored at −80°C until processing for HPLC. As previously described, tissue was thawed on ice and sonicated in 0.1 N perchloric acid (10 μl/mg tissue) for 12 s with 0.5 s pulses. Sonicated samples were centrifuged (16,100 rcf) for 30 min at 4 °C, and the supernatant was then centrifuged through 0.45 μm filters at 4000 rcf for 10 min at 4 °C.

For HPLC, an ESA 5600A CoulArray detection system equipped with an ESA Model 584 pump and an ESA 542 refrigerated autosampler was used. Separations were performed at 28 °C using an MD-150 × 3.2 mm C18 column. The mobile phase consisted of 1.6 mM 1-octanesulfonic acid sodium, 75 mM NaH2PO4, 0.025% triethylamine, and 8% acetonitrile at pH 2.98. 20 μl of sample was injected. The samples were eluted in an isocratic manner at 0.4 ml/min and detected using a 6210 electrochemical cell (ESA, Bedford, MA) equipped with 5020 guard cell. Guard cell potential was set at 475 mV, while analytical cell potentials were − 175, 150, 350 and 425 mV. NE and DA were measured by electrochemical detection. The analytes were identified by the matching criteria of retention time to known standards (Sigma-Aldrich). Compounds were quantified by comparing peak areas to those of standards on the dominant sensor.

### 2.4 Immunohistochemistry and Fluorescent Imaging

The tissue used for the detection of the NE transporter (NET), tyrosine hydroxylase (TH; total TH), and phospho-Ser31 TH by immunohistochemistry (IHC) was collected from an experiment in which *Dbh+/−* control mice were exposed to odorless cage change or cage change in the presence of the novel odorant (n = 3 male mice per condition, 8 months of age). Mice were euthanized 30 min after exposure to either odorless cage change or cage change with a novel odorant (palmarosa) with an overdose of sodium pentobarbital (Fatal Plus, 150 mg/kg, i.p.; Med-Vet International, Mettawa, IL) prior to perfusion with cold 4% paraformaldehyde/0.01 M PBS buffer. The tissue used for the detection of DBH in the LC of *Dbh*+/− and *Dbh*−/− mice came from experimentally naïve animals (n = 3 male mice per genotype, 8 months of age).

Following extraction, brains were post-fixed for 24 h in 4% paraformaldehyde/0.01 M PBS buffer at 4°C, and then transferred to cryoprotectant 30% sucrose/ 0.01M PBS solution for 72 h at 4°C. Brains were embedded in OCT medium (Tissue-Tek; Sakura, Torrance, CA) and serially sectioned by cryostat (Leica, Wetzlar, Germany) into 40-μm coronal slices containing the LC and PFC. Brain sections were stored in 0.01 M PBS buffer (+ 0.02% sodium azide) at 4°C before IHC.

For IHC, brain sections were blocked for 1 h at room temperature in 5% normal goat serum (NGS; Vector Laboratories, Burlingame, CA) diluted in 0.01 M PBS/0.1% Triton-X permeabilization buffer. Sections were then incubated for 24 h at 4°C in NGS blocking and permeabilization buffer, including primary antibodies raised against DBH (rabbit anti-DBH; Immunostar, Hudson, WI, #22806; 1:1000), the NE transporter (NET) (mouse anti-NET; MAb Technologies, Neenah, WI, NET05-2; 1:1000), total TH (rabbit anti-TH; P40101–0, Pel-Freez, Rogers, AR; 1:1000), and phospho-Ser31 TH (rabbit anti-phosphoSer31TH; Cell Signaling Technology, Danvers, MA, #13041; 1:1000). After washing in 0.01 M PBS, sections were incubated for 2 h in blocking/permeabilization buffer with goat anti-mouse and/or goat anti-rabbit AlexaFluor 568 (Invitrogen, Carlsbad, CA; 1:500) secondary antibodies. After washing again, the sections were mounted onto Superfrost Plus slides (Thermo Fisher, Waltham, MA) and cover-slipped with Fluoromount-G plus DAPI (Southern Biotech, Birmingham, AL).

Fluorescent micrographs of immunostained sections were acquired on a Leica DM6000B epifluorescent upright microscope at 20× magnification with uniform exposure parameters. For the quantification experiments measuring LC and PFC expression of total TH and phospho-Ser31 TH, micrographs were acquired at 20× for visualization of LC cell bodies and axon terminals in the prelimbic cortex (PrL), a subregion of the medial PFC (mPFC). We selected atlas-matched sections from each animal at the level of the LC and mPFC. A standardized region of interest was drawn for all images to delineate the borders of the LC and PrL in all animals. Image processing and analysis were performed using ImageJ software. The analysis pipeline included standardized background subtraction, intensity thresholding (Otsu method), and pixel intensity measurements within defined regions of interest cropped to the same size, as previously described (Foster et al., 2021; Lustberg et al., 2020b).

### 2.5 Statistical Analysis and Figure Design

For the odorless cage change (+ water) or novel odorant cage change (+ tea tree oil) experiment, the effects of odorant on repetitive behaviors in *Dbh*+/− and *Dbh*−/− mice were compared using two-way repeated measures ANOVAs (genotype x odorant), with post hoc Bonferroni tests for multiple comparisons computed within genotypes and between odorants. The effects of anti-adrenergic drugs on repetitive behaviors compared to saline vehicle were assessed using a one-way ANOVA for treatment, with post hoc Dunnett’s test for multiple comparisons. The effects of flupenthixol on repetitive behaviors in *Dbh*−/− mice were compared using a paired sample t-test.

The effects of odorant on plasma CORT levels and central catecholamine (NE and DA) abundance were compared within the dorsal pons/LC and PFC between *Dbh* genotypes and across odorless cage change and cage change + novel odorant conditions by two-way ANOVAs (genotype x odorant), with post hoc Bonferroni tests for multiple comparisons performed within genotypes and between odorant conditions.

In *Dbh+/−* control mice treated with either saline or nepicastat prior to novel odorant exposure, pons and PFC samples were compared for levels of central catecholamines using multiple t-tests with Holm-Sidak corrections for multiple comparisons. Plasma CORT levels were compared between saline and nepicastat treatment groups using an unpaired sample t-test.

In *Dbh+/−* control mice euthanized 30 min after odorless cage change or cage change with a novel odorant, total TH and phospho-Ser31 TH immunoreactivity in the LC and PFC were compared within regions and between conditions using multiple t-tests, with Holm-Sidak corrections for multiple comparisons.

The threshold for adjusted significance was set at p < 0.05, and two-tailed variants of tests were used throughout. Analyses and graph design were performed using Prism v9 (GraphPad Software, San Diego, CA). Where appropriate, effect sizes (Cohen’s d) were computed using this free resource: https://www.psychometrica.de/effect_size.html.

## 3. Results

### 3.1 DBH deficiency suppresses odorant-induced defensive digging while facilitating grooming

We assessed two subtypes of digging behavior, as well as grooming, when *Dbh−/−* and control mice were transferred to a new clean cage and exposed to either water (odorless control) or tea tree oil (novel odorant). “Light digging” was defined as coordinated movement of the front or back paws which displaced a moderate amount of bedding while the mouse was otherwise stationary (De Boer and Koolhaas, 2003; Pond et al., 2021). Light digging is considered exploratory in nature because it occurs broadly across a range of experimental conditions if a bedding substrate is provided (Kemble and Bolwahnn, 1997; Londei et al., 1998; Wolmarans et al., 2016).

By contrast, “defensive burying” involved vigorous displacement of the bedding in the direction of the novel odorant stimulus, often resulting in the burying of the odorant-soaked nestlet. Defensive burying sometimes presented as intense burrowing away from the stimulus, which resembled “swimming” or tunneling through the bedding substrate (Bondi et al., 2007; De Boer and Koolhaas, 2003; Fucich and Morilak, 2018; Treit et al., 1981). Burying is considered an aversion-related responses because it resembles responses exhibited after exposure to the shock probe test or predator odor (Garbe et al., 1993; Hwa et al., 2019; McGregor et al., 2002; Sluyter et al., 1996).

Grooming was defined as repetitive paw movements oriented around the whiskers and face, as well as licking or scratching of the body and tail (De Boer and Koolhaas, 2003; Kalueff et al., 2016; Smolinsky et al., 2009). As with digging, some amount of grooming is normal for mice and is a marker of wellbeing, but the behavioral link between stress and grooming is less clear (De Boer and Koolhaas, 2003; Guild and Dunn, 1982; Kalueff and Tuohimaa, 2005; Smolinsky et al., 2009). For instance, some reports suggest that grooming may be a self-soothing behavior that increases under stressful conditions, while others propose that it is a “rest behavior’ that indicates low levels of stress (Guild and Dunn, 1982; Kemble and Bolwahnn, 1997; Kemble et al., 1995).

There was a strong trend for reduced “light digging” behavior in the *Dbh−/−* mice compared to controls that did not reach statistical significance [F(1,11) = 4.75, p = 0.052], but no main effect of odorant [F(1,11) = 0.84, p = 0.38] or odorant x genotype interaction [F(1,11) = 0.45, p = 0.52] (Fig. 1A). However, when we compared defensive burying between genotypes and odorant conditions, we found an overall main effect of genotype [F(1,11) = 6.83, p = 0.02, d = 1.58], a main effect of odorant [F(1,11) = 8.52, p = 0.01, d = 1.77], and a genotype x odorant interaction [F(1,11) = 6.83, p = 0.02, d = 1.58] (Fig. 1B). NE-competent control mice only engaged in defensive burying behavior in response to the novel odorant stimulus following cage change (p < 0.01 compared to water as the odorless control condition), while *Dbh−/−* mice virtually never engaged in defensive burying after cage change irrespective of the novel odorant (p > 0.99) (Vid. 1).

**Fig. 1.**
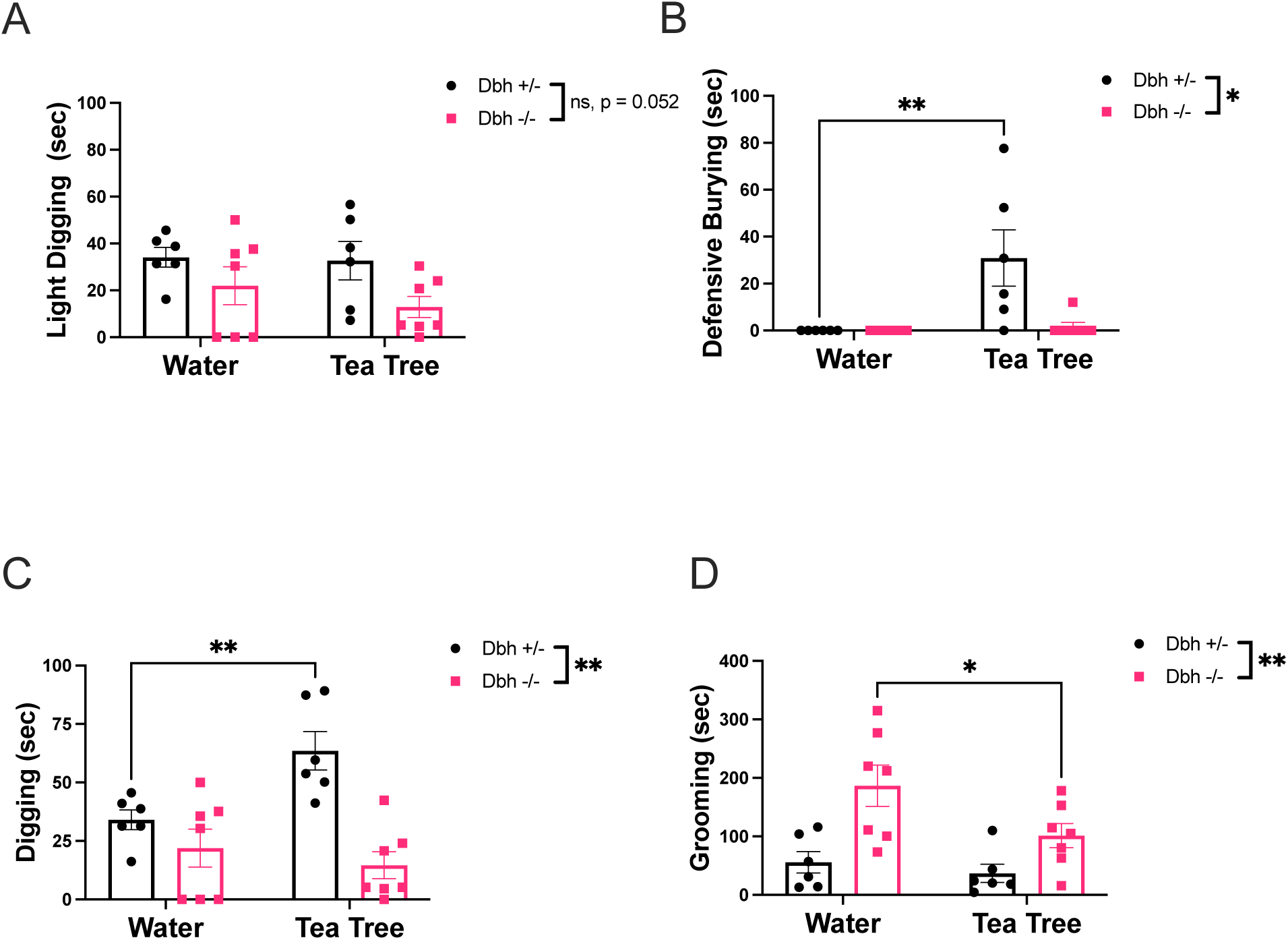
Effect of cage change with novel odorant (tea tree) compared to odorless water on task duration spent (sec) engaged in light digging **(A)** and defensive burying **(B)**, which together comprise a composite score for time spent engaged in total digging for mice of both genotypes **(C)**. Mice of both *Dbh* genotypes were also assessed for time spent grooming **(D)**. (*) and (**) indicate p < 0.05 and p < 0.01, respectively. (ns) indicates p > 0.05. Error bars denote ±SEM.

**Vid. 1.**
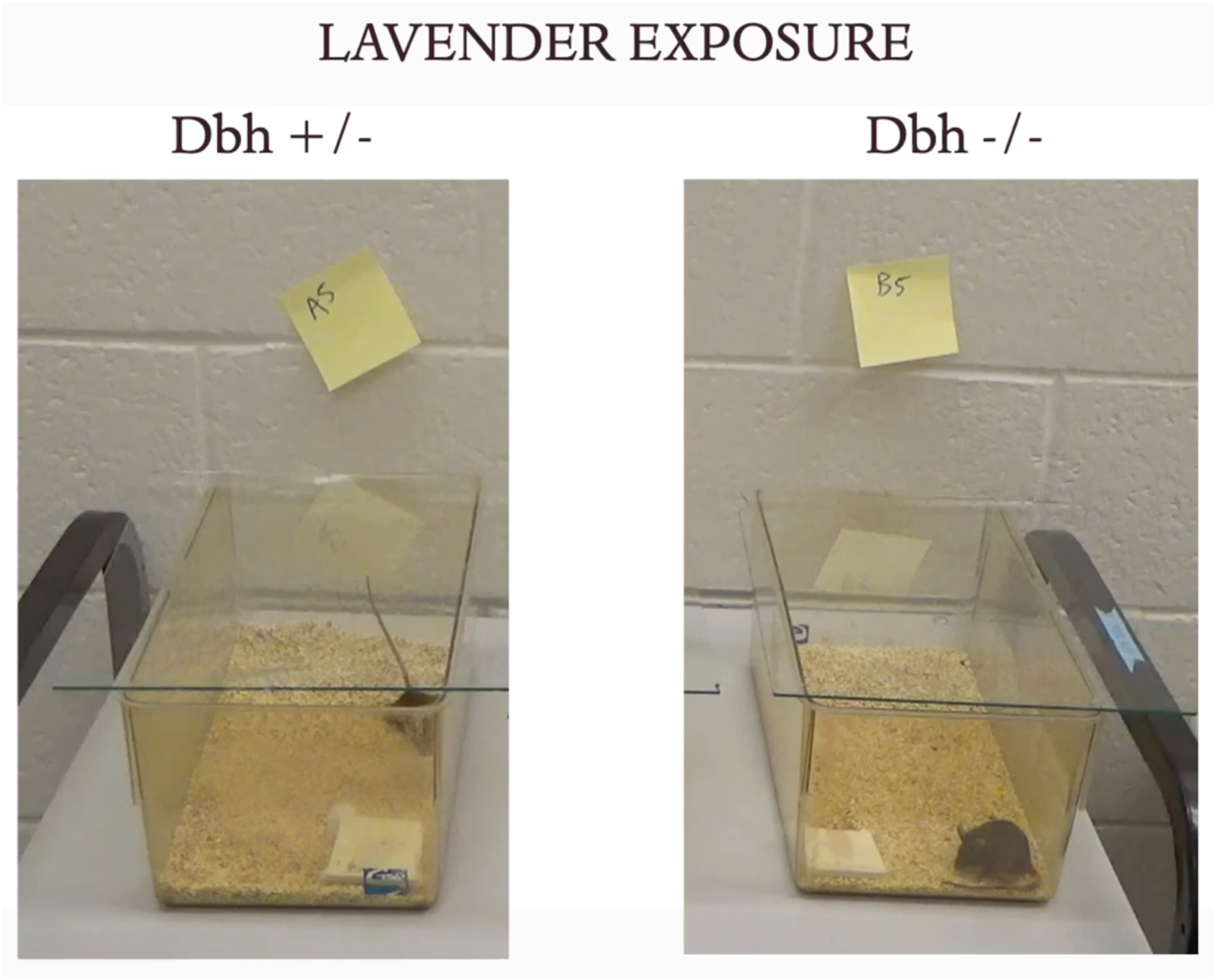
Simultaneous recording of *Dbh+/−* (left) and *Dbh−/−* (right) mice exposed to a new clean cage scented with lavender oil as a novel odorant.

Given that “light digging” and defensive burying are both subtypes of overall digging behavior, and that the genotype difference in cage change-induced light digging nearly reached statistical significance between genotypes, we also calculated a composite digging score for each genotype and odorant condition. There was an overall main effect of genotype [F(1,11) = 14.28, p < 0.01, d = 2.29] and a genotype x odorant interaction [F(1,11) = 11.86, p < 0.01, d = 2.08], as well as a trend for a main effect of odorant that did not reach statistical significance [F(1,11) = 4.29, p = 0.06] (Fig. 1C). Post-hoc analysis revealed that *Dbh+/−* mice demonstrated more overall digging in the novel odorant condition compared to the odorless control condition (p < 0.01), while the digging behavior of *Dbh−/−* mice did not differ as a function of odorant (p = 0.67).

When we assessed grooming, a non-defensive repetitive behavior, we found an overall main effect of genotype [F(1,11) = 12.94, p < 0.01, d = 2.18] and odorant [F(1,11) = 5.51, p = 0.04, d = 1.42], but no genotype x odorant interaction [F(1,11) = 2.24, p = 0.16] (Fig. 1D). *Dbh−/−* mice groomed significantly more than *Dbh+/−* control mice irrespective of odorant condition but groomed less frequently in the novel odorant condition than in the odorless cage change condition (p = 0.03) (Vid. 1). Control mice did not display differences in grooming behavior between novel odorant conditions (p > 0.99).

### 3.2 Pharmacological blockade of NE transmission reduces stress-induced digging behavior in control mice, while DA signaling receptor antagonism suppresses excessive grooming in Dbh−/− mice

We assessed the effects of drugs that block the NE synthesis, release, or AR signaling on the duration of novel odorant-induced overall digging in *Dbh+/−* control mice, as a composite of “light digging” and “defensive burying.” Although grooming did not differ as a function of novel odorant exposure in *Dbh+/−* control mice in our previous experiments, we also measured grooming to account for any non-specific drug effects on motor behavior. The drugs tested included the DBH inhibitor nepicastat to mimic genetic DBH knockout, the inhibitory α_2_-autoreceptor agonist guanfacine to suppress NE release, and a cocktail of the β_1/2_AR antagonist propranolol and α_1_AR antagonist prazosin to block postsynaptic NE signaling. We found a main effect of drug treatment compared to saline treatment [F(3,43) = 15.19, p < 0.0001, d = 1.17] (Fig. 2A). Overall digging behavior was potently suppressed by nepicastat (p < 0.01), guanfacine (p < 0.0001), or the cocktail of AR antagonists (p < 0.0001). These results indicate that the abnormally infrequent digging observed in the *Dbh−/−* mice reflects an acute lack of NE at the time of the test rather than compensation in other systems in response to lifelong NE deficiency.

**Fig. 2.**
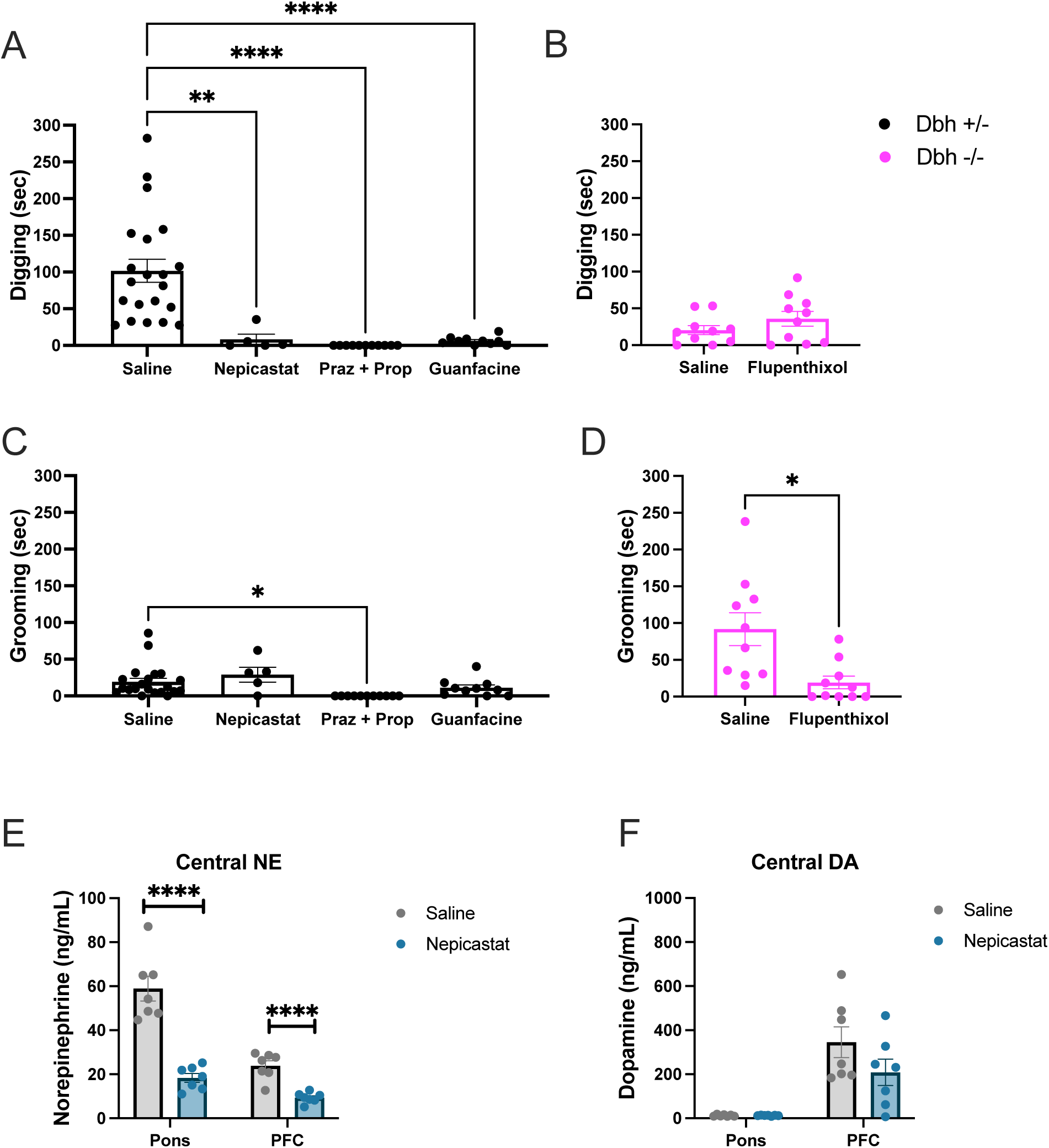
Effects of anti-adrenergic drugs on novel odorant-induced digging **(A)** and grooming **(C)** behaviors in *Dbh+/−* mice, and of the DA receptor antagonist flupenthixol on the same behaviors in *Dbh−/−* mice **(B, D)**. Effects of nepicastat treatment on NE **(E)** and DA **(F)**levels in the dorsal pons/LC and PFC of *Dbh+/−* mice exposed to cage change with a novel odorant. (*), (**), and (****) indicate p < 0.05, p < 0.01, and p < 0.0001 compared with the vehicle-treated control group, respectively. Error bars denote ±SEM.

For grooming, there was a main effect of drug treatment [F(3,43) = 4.46, p < 0.01, d = 0.63] (Fig. 2C). Posthoc analysis revealed that the cocktail of NE receptor antagonists prazosin and propranolol significantly reduced grooming (p = 0.01), while the other compounds (nepicastat, p = 0.58; guanfacine, p = 0.51) had no effect.

The cocktail of prazosin and propranolol reduced both digging and grooming in control mice, and in fact this drug combination resulted in profound cessation of all observable behaviors in the test environment. Despite inducing behavioral arrest, the cocktail of AR antagonists did not cause noticeable sedation or loss of coordination, and normal motor behavior returned immediately upon placement back in the home cage. These results suggest a unique interaction between cage change + novel odorant stress and simultaneous blockade of β- and α1 ARs.

Because treatments that targeted NE signaling attenuated digging but did not increase grooming, we suspected that the aberrant grooming we observed in *Dbh−/−* mice was driven by the supranormal levels of DA in these animals (Bourdélat-Parks et al., 2005; Thomas et al., 1998; Weinshenker et al., 2002a). In *Dbh−/−* mice, pre-treatment with the DA receptor antagonist flupenthixol had no effect on digging [t(9) = 1.22, p = 0.25] (Fig. 2B), but significantly reduced grooming behavior in the presence of a novel odorant compared to saline treatment [t(9) = 2.82, p = 0.02, d = 1.40] (Fig. 2D).

Given that excessive grooming in the *Dbh−/−* mice appeared to be driven by DA, and we and others have reported that nepicastat increases tissue levels of DA in normal animals (Acosta et al., 2022; Devoto et al., 2015; Schroeder et al., 2010), it was puzzling that nepicastat did not increase grooming in *Dbh+/−* mice. To investigate potential neurochemical mechanisms, we assessed the effects of nepicastat on catecholamine levels. Consistent with its ability to suppress digging, nepicastat reduced NE content in the dorsal pons/LC [t(12) = 6.80, p < 0.0001, d = 3.64] and PFC [t(12) = 5.94, p < 0.0001, d = 3.18] (Fig. 2E), as expected, but had no effect on DA levels in pons [t(12) = 0.12, p = 0.91] or PFC [t(12) = 1.48, p = 0.39]) compared to saline (Fig. 2F). These results confirm that digging is primarily mediated by NE and provide an explanation for the inability of nepicastat to increase grooming in *Dbh+/−* mice.

### 3.3 Novel odorant exposure induces regional changes in central catecholamine content

Stress activates the LC, and NE and DA are released in LC targets like the PFC during stress to facilitate adaptive and flexible behaviors critical for survival (Arnsten et al., 2017; Aston-Jones et al., 1999; McCall et al., 2015; Ross and Van Bockstaele, 2021). Given the link between catecholamine signaling and stress reactivity, we measured central levels of NE and DA in the dorsal pons/LC and PFC in mice of both *Dbh* genotypes following transfer to a new clean cage (odorless control condition) or a new clean cage with a novel odorant (palmarosa). We found the expected main effect of *Dbh* genotype [F(1,26) = 188.10, p < 0.0001, d= 4.12] on NE levels in the pons (Thomas et al., 1998), but no effect of odorant [F(1,26) = 0.38, p = 0.54] or genotype x odorant interaction [F(1,26) = 0.38, p = 0.54] (Fig. 3A). These results confirm that *Dbh−/−* mice lack pontine NE but suggest that novel odorant exposure does not increase brainstem NE levels above those induced by odorless cage change in control mice.

**Fig. 3.**
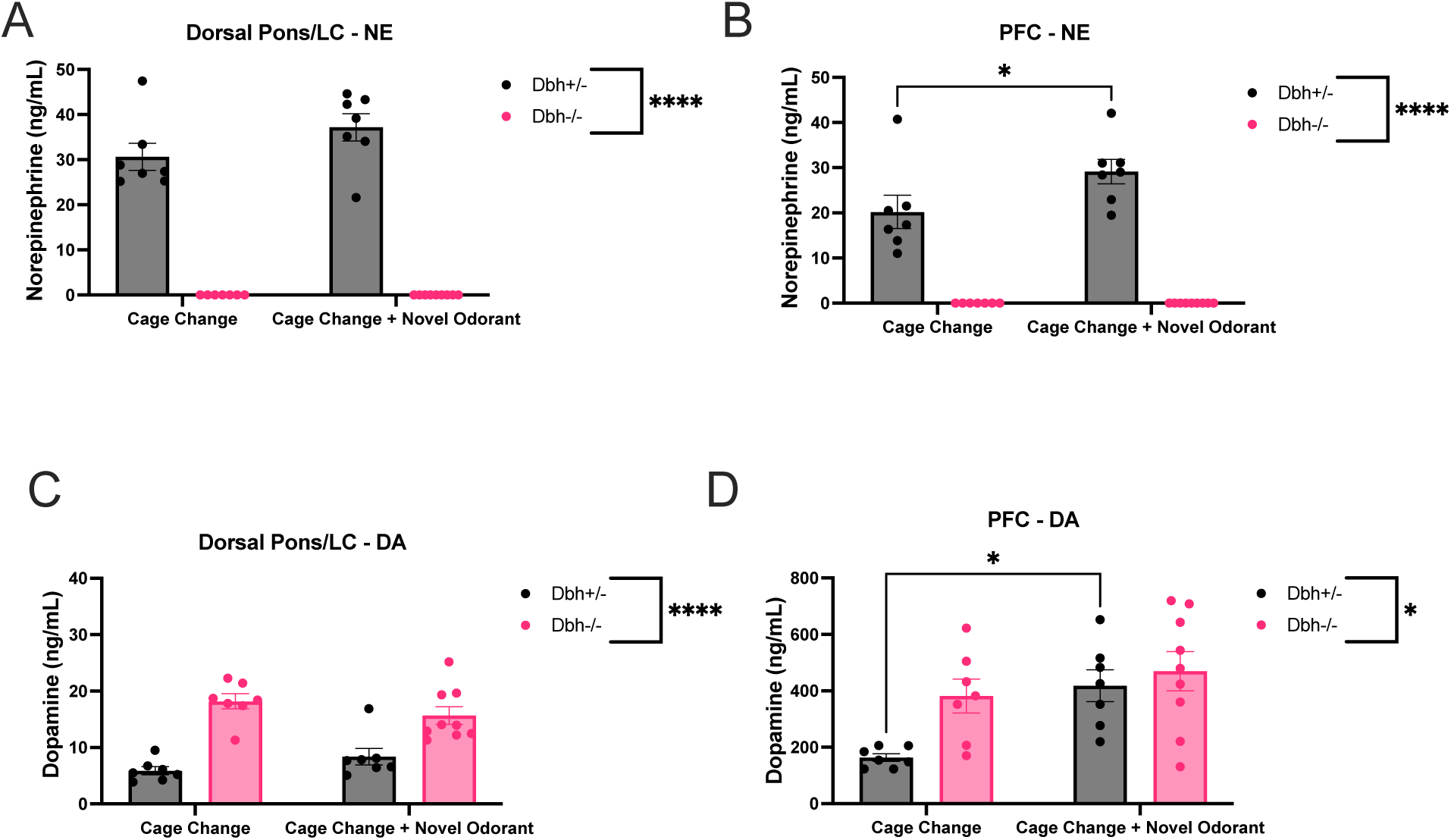
Effects of cage change with or without a novel odorant on NE levels in the dorsal pons/LC **(A)** and PFC **(B)** of *Dbh−/−* and *Dbh+/−* mice, as well as effects of cage change with or without a novel odorant on DA levels in the dorsal pons/LC (**C)** and PFC **(D)** of *Dbh−/−* and *Dbh+/−* mice. (*) and (****) indicate p < 0.05 and p < 0.0001, respectively. Error bars denote ±SEM.

In the PFC, there was a main effect of genotype [F(1,26) = 133.70, p < 0.0001, d= 4.38], odorant [F(1,26) = 4.41, p = 0.04, d= 0.80], and a genotype x odorant interaction [F(1,26) = 4.41, p = 0.04, d= 0.80] for tissue levels of NE (Fig. 3B). In contrast to the dorsal pons/LC, novel odorant exposure increased PFC NE content in control mice (p = 0.02) but not *Dbh−/−* mice (p > 0.99) compared to the odorless cage change condition. Thus, cage change with a novel odorant exposure increases PFC levels of NE in control mice compared to cage change in the absence of a novel odorant.

Although some of the DA in the PFC originates from the VTA (Kiyatkin, 1995; Schmidt et al., 2017), LC terminals in the PFC also contain and release DA (Devoto et al., 2015; Devoto et al., 2020). Thus, we measured DA in the dorsal pons/LC and PFC of *Dbh+/−* and *Dbh−/−* mice following transfer to a new clean cage (odorless control) or a new clean cage with a novel odorant (palmarosa). In line with previous reports, we found a main effect of genotype [F(1,26) = 51.15, p < 0.0001, d = 2.71] such that *Dbh−/−* mice had higher levels of pontine DA than control mice (Thomas et al., 1998), but there was no effect of odorant [F(1,26) < 0.001, p > 0.99] and a non-significant trend for a genotype x odorant interaction [F(1,26) = 3.34, p = 0.08] (Fig. 3C). These results confirm that *Dbh−/−* mice have excessive levels of pontine DA compared to control mice but suggest that novel odorant exposure does not increase pontine DA levels above those induced by odorless cage change in either control or *Dbh−/−* mice.

In the PFC, there was main effect of genotype [F(1,26) = 5.50, p = 0.03, d = 0.88] and odorant [F(1,26) = 8.87, p < 0.01, d = 1.12] for tissue levels of DA, but no genotype x odorant interaction [F(1,26) = 2.10, p = 0.16] (Fig. 3D). In contrast to the dorsal pons/LC, odorant exposure increased PFC levels of DA in control mice (p = 0.01) but not *Dbh−/−* mice (p = 0.55), which were higher than those of *Dbh+/−* control mice regardless of odorant condition. Taken together, these results confirm that *Dbh−/−* mice have excessive levels of PFC DA and reveal that olfactory novelty has brain region- and genotype-specific effects on DA. The ability of DBH knockout, but not nepicastat, to increase DA levels in the presence of cage change + novel odorant stress may also explain why the former but not the latter manipulation induces robust grooming behavior under these experimental conditions.

### 3.4 Novel odorant exposure induces tyrosine hydroxylase phosphorylation in the PFC

Fluorescence immunohistochemistry (IHC) of the LC of *Dbh+/−* and *Dbh−/−* mice confirmed that the gross structure of the LC remains intact despite the total lack of DBH protein (Jin et al., 2004) (cyan; Fig. 4a). Notably, LC neurons of *Dbh−/−* mice still express NET (red; Fig. 4A) (Jin et al., 2004; Sanders et al., 2006; Weinshenker et al., 2002b).

**Fig. 4.**
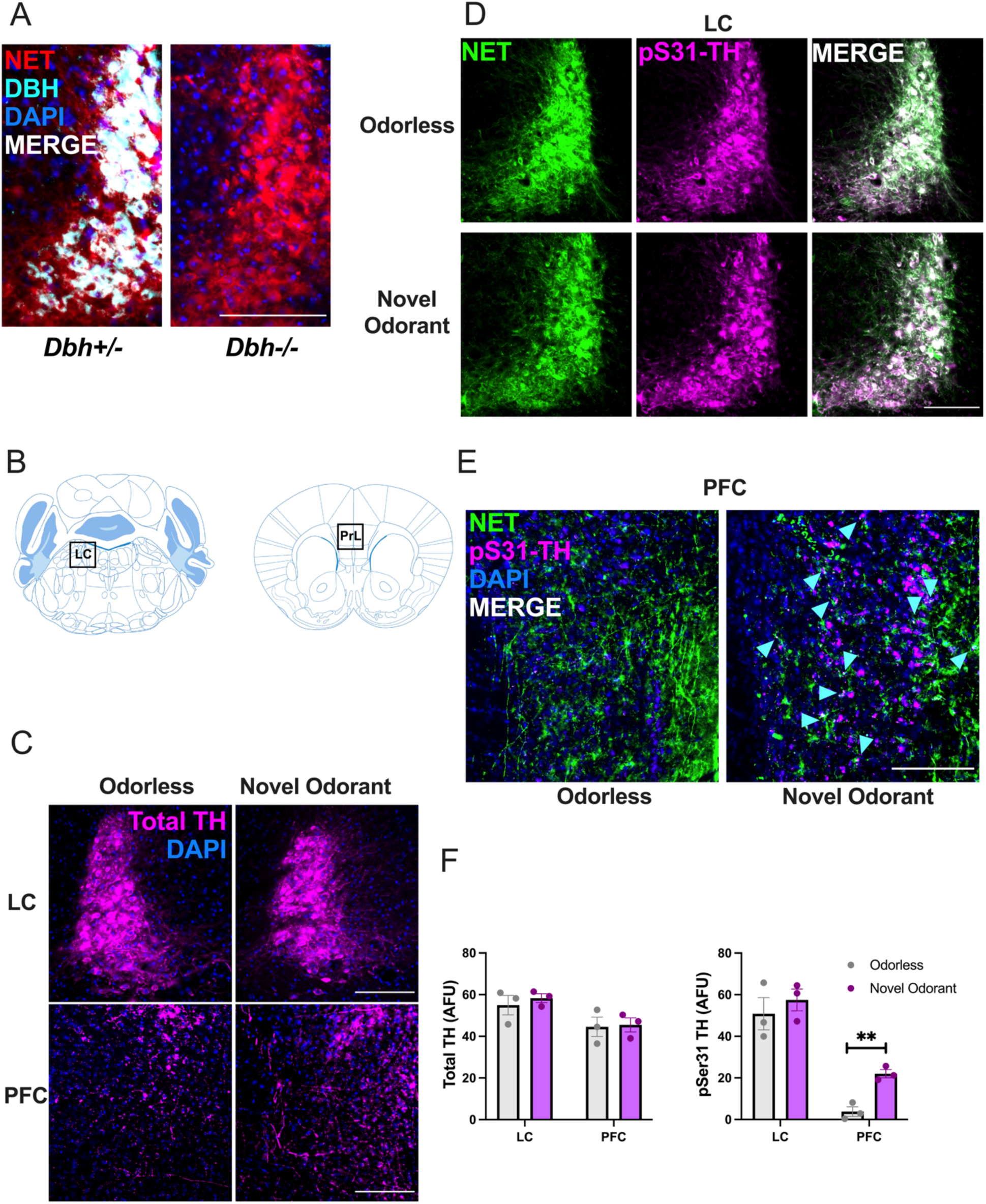
Effects of novel odorant exposure on TH phosphorylation in the LC and PFC **(B)**. *Dbh−/−* mice lack DBH (cyan) but not NET (red) expression in the LC **(A)**. Total TH immunoreactivity (magenta) in arbitrary fluorescent units (AFUs) in the LC and the PrL subregion of the PFC **(C, F)** of *Dbh+/−* mice exposed to either odorless cage change or cage change with a novel odorant. Immunoreactivity for phospho-Ser31 TH (magenta) in the LC (**D, F)** and PFC (**E, F)** of the same animals. In (**E)**, cyan arrows denote patches where NET (green) and phospho-Ser31 TH overlap. (**) indicates p < 0.01, and scale bars in micrographs denote 100 μm. Error bars denote ±SEM.

Because we detected an increase in PFC concentrations of both catecholamines 30 min after exposure to a novel odorant, we next investigated whether novel odorant exposure altered the phosphorylation state of the enzyme tyrosine hydroxylase (TH) (Daubner et al., 2011). Phosphorylation of TH, which converts tyrosine into L-DOPA and is the rate-limiting enzyme in the synthesis of both DA and NE, rapidly increases the enzymatic rate of catecholamine production (Dunkley and Dickson, 2019; Ong et al., 2014).

Using *Dbh+/−* control mice exposed to either odorless cage change or cage change with a novel odorant (palmarosa), we measured expression of TH phosphorylated at serine reside 31 (phospho-Ser31 TH) as well as total TH in the LC and PrL subregion of the PFC 30 min after each exposure (Fig. 4B). We found that novel odorant exposure increased expression of phospho-Ser31 TH in the PFC [t(4) = 6.03, p < 0.01, d = 4.9] but not the LC [t(4) = 0.71, p = 0.51] compared to odorless cage change (Fig. 4D-F). There was no effect of novel odorant exposure on total TH levels in the LC [t(4) = 0.66, p = 0.55] or PFC [t(4) = 0.16, p = 0.88] compared to odorless cage change (Fig. 4C,F), suggesting that the increase in catecholamine levels elicited by the novel odorant exposure may be explained by rapid local post-translational modification (phosphorylation) of TH at residue Ser31 in terminals but not cell bodies/proximal dendrites (Jorge-Finnigan et al., 2017; Ong et al., 2014; Ong et al., 2011; Salvatore et al., 2001).

### 3.5 Plasma corticosterone (CORT) levels do not differ as a function of odorant condition or catecholamine status

The lipophilic hormone CORT is synthesized in the adrenal gland and is the major endocrine mediator of the physiological stress response in rodents (de Quervain et al., 2017; McKlveen et al., 2015). An elevation in circulating plasma CORT levels is the “gold standard” physiological correlate of stress in animal models (Bowers et al., 2008; Mora et al., 2012). Because cage change with a novel odorant elicited behavioral and neurochemical responses reflective of increased stress reactivity compared to cage change alone and depended on activity, we hypothesized that novel odorant exposure would increase blood levels of CORT, with potential differences in mice lacking NE.

We measured plasma CORT abundance following transfer to a new clean cage (odorless control) or a new clean cage with a novel odorant (palmarosa) in *Dbh+/−* and *Dbh−/−* mice. There were no main effects of genotype [F(1,20) = 1.43, p = 0.25] or odorant [F(1,20) = 0.12, p = 0.72] on CORT levels, and no genotype x odorant interaction [F(1,20) = 0.36, p = 0.55] (Fig. 5A), suggesting that novel odorant exposure did not elicit a systemic stress response above and beyond cage change. There was also no effect of nepicastat pre-treamtent on CORT levels induced by novel odorant stress compared to saline [t(11) = 0.42, p = 0.68] (Fig. 5B). These results suggest that plasma CORT sampled at this time point (30 min) is a relatively insensitive measure of psychological stress induced by novelty and does not reflect different stress-induced behavioral sequelae as clearly as changes in PFC catecholamines (Finlay et al., 1995; Jimeno et al., 2018; MacDougall-Shackleton et al., 2019).

**Fig. 5.**
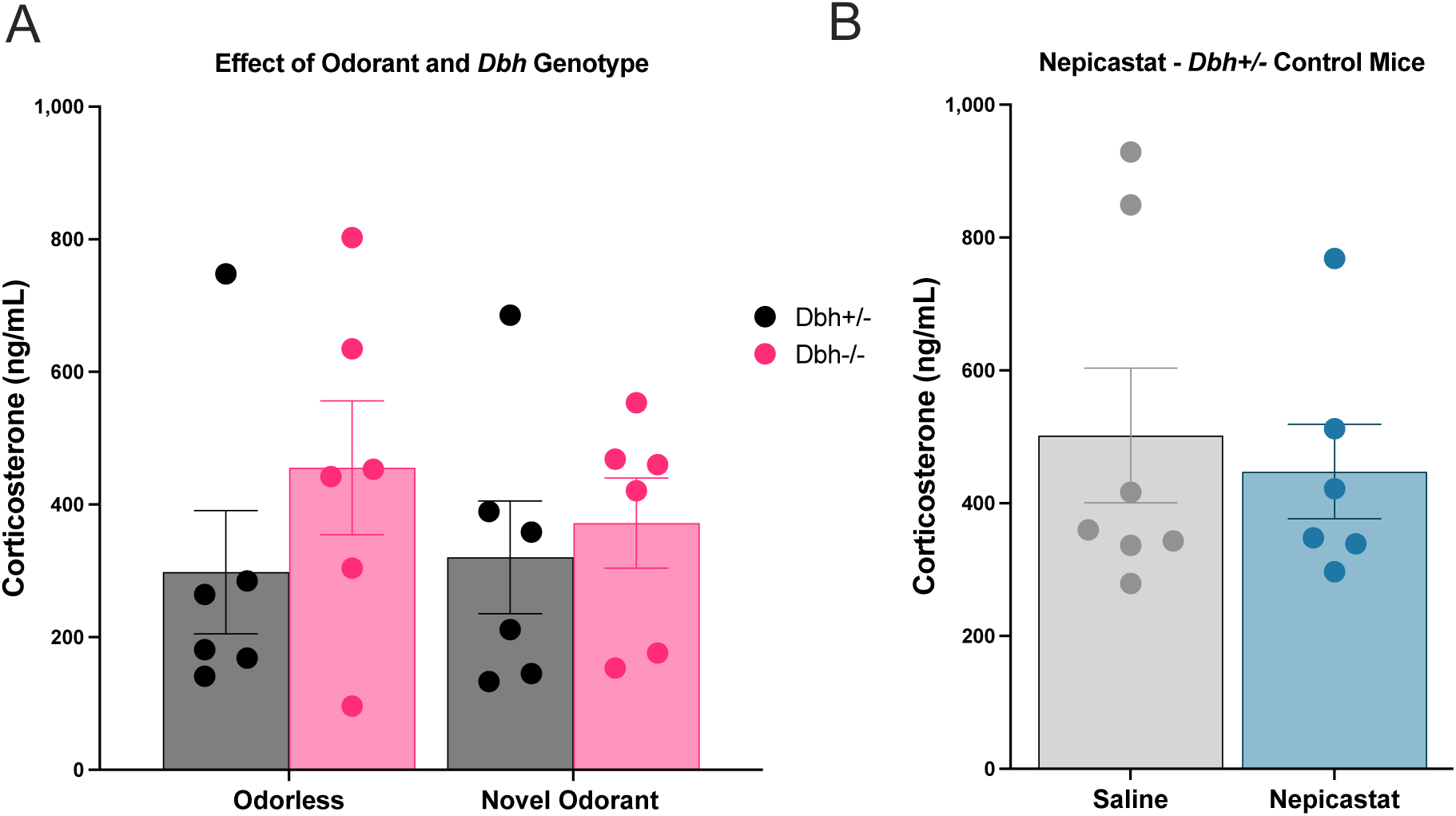
Effects of genetic **(A)** or pharmacological **(B)** depletion of NE on plasma CORT levels following cage change with or without a novel odorant. All mice in the nepicastat treatment experiment were *Dbh+/−* control mice exposed to cage change with a novel odorant. Error bars denote ±SEM.

## 4. Discussion

We assessed the effects of manipulating catecholamine transmission on innate, repetitive behaviors induced by novel odorant stress using *Dbh−/−* mice and their NE-competent littermate controls, as well as pharmacological tools. Further, we measured changes in brain concentrations of catecholamines and the phosphorylation status of the catecholamine synthetic enzyme TH in the LC and PFC of *Dbh−/−* and *Dbh+/−* control mice following exposure to novel plant odorants, as well as plasma CORT levels. The pharmacological approaches of this study clarify certain experimental findings which might otherwise be confounded by compensatory adaptations or developmental differences associated with life-long, global loss of DBH function in *Dbh−/−* mice.

Confrontation with a novel environment or stimulus elicits a stress response in rodents (Garbe et al., 1993; Masuo et al., 2021; Seggie and Brown, 1975), which mobilizes cognitive and metabolic resources required for adaptive behavior under conditions of uncertainty or change (Aston-Jones et al., 1999). Systemic endocrine (e.g. CORT) and central neuromodulator (e.g. catecholamine) levels are augmented by stress (Irwin et al., 1986), which can acutely alter cognitive and affective responses in rodents through actions on forebrain networks that govern behavioral reactivity (McKlveen et al., 2015; Mora et al., 2012; Ross and Van Bockstaele, 2021).

Digging, encompassing both exploratory digging (low-stress) and defensive burying (high-stress), is a stress-sensitive repetitive behavior observed across a range of behavioral tests in which a bedding substrate is provided, including the shock probe and MB tests (De Boer and Koolhaas, 2003; Hwa et al., 2019; Wolmarans et al., 2016). In the shock probe test, burying is considered an active defense behavior to cope with the physical stressor of the shock delivered by the probe (Fucich and Morilak, 2018; Sluyter et al., 1996). Grooming, although less clearly elicited by stress than digging, is a repetitive behavior that can be triggered by psychological stressors or even central infusions of the pituitary stress hormone ACTH (Guild and Dunn, 1982; Smolinsky et al., 2009).

In our study, a clean cage scented with novel plant-derived odorant (tea tree oil) evoked more defensive burying in control mice compared to a clean cage treated with odorless water. In the shock probe task, defensive burying is interpreted as an active coping response to a stressor as opposed to the passive coping response of freezing (Bondi et al., 2007; Fucich and Morilak, 2018). In the MB test, which measures digging but does not classically discriminate between digging and defensive burying, the marbles are buried as consequence of overall digging behavior in the substrate (de Brouwer et al., 2019; Dixit et al., 2020). Digging in the MB test is frequently interpreted a repetitive, compulsive-like behavior, although some researchers consider it a model of anxiety-like behavior (de Brouwer et al., 2019; Londei et al., 1998). Here, we found that *Dbh* deficiency or drugs that block NE synthesis or transmission suppress novel odorant-induced digging behavior, and we have previously shown that *Dbh* deficiency or the same battery of drugs tested in this study reduce nestlet shredding and marble burying in other assays of stress-induced repetitive behavior (Lustberg et al., 2020a). Combined, these results bolster support for the hypothesis that central NE is required for stress-induced repetitive digging (Bondi et al., 2007; Howard et al., 2008; Lustberg et al., 2020a).

Grooming is not classically considered to be a coping behavior in response to a stressor; animals will self-groom under low and/or high stress conditions (Kalueff et al., 2016; Smolinsky et al., 2009). However, animal models of obsessive-compulsive disorder (OCD) and Tourette Syndrome (TS) demonstrate high levels of repetitive self-grooming behavior (Manning et al., 2021; Nordstrom and Burton, 2002; Welch et al., 2007), and grooming can be elicited by central infusions of ACTH or DA agonists (Guild and Dunn, 1982; Taylor et al., 2010). The ACTH-induced grooming response was prevented by drugs that block DA receptors, while grooming induced by central ACTH infusion is augmented by the DA agonist apomorphine (Guild and Dunn, 1982). Administration of DA agonists elicit exaggerated grooming behavior, and constitutive D1R activation in the D1CT7 transgenic mouse generates a TS-like phenotype characterized by excessive grooming and tics (Godar et al., 2016; Nordstrom et al., 2015). These results implicate abnormal DA signaling in excessive grooming behavior (Taylor et al., 2010; Wood et al., 2018).

We found that, regardless of odorant, *Dbh−/−* mice exhibited frequent and extended bouts of repetitive grooming behavior, which was rarely observed in control mice regardless of odorant condition. Moreover, the nonspecific DA receptor antagonist flupenthixol reduced abnormal grooming in *Dbh−/−* mice without altering digging behavior. Thus, excessive DAergic signaling in the *Dbh−/−* mice appears to be caused by increased central DA signaling or DA receptor adaptations in *Dbh* mutants (Sanders et al., 2006; Schank et al., 2006; Weinshenker et al., 2002a).

Our results indicate that NE governs stress-induced digging, while DA drives grooming. These distinct repetitive behaviors are remarkably dependent on distinct types of catecholaminergic signaling. That said, a few exceptions in our results are worth noting. The unique suppressive effect of blocking β_1/2_ARs and α_1_ARs on both digging and grooming may be explained by a peculiar behavioral phenomenon that occurred only in the novel odorant exposure condition in mice treated with the AR antagonist cocktail. Mice administered the prazosin and propranolol cocktail demonstrated complete behavioral arrest when transferred to the novel-scented cage, involving the abolition of all observable exploratory, locomotor, or repetitive behavior. Although virtually motionless, mice treated with the AR antagonist cocktail retained consciousness, maintaining upright posture and displaying at least partial responsiveness to external stimuli. For example, the mice attempted to avoid being picked up by the tail, and the behavioral arrest ceased immediately upon return of the animal to the home cage environment. These pharmacological findings, though unexpectedly contingent upon the presence of a novel odorant in the test environment, are consistent with previous reports in which central infusion of α_1_ AR antagonists induce cataleptic-like responses following cage change (Lin et al., 2008; Stone et al., 2001; Stone et al., 2006; Stone et al., 1999).

Another intriguing finding was that the DBH inhibitor nepicastat recapitulated the reduced digging observed in *Dbh−/−* mice but failed to similarly elicit excessive grooming. Given that NE is permissive for digging while high DA drives grooming, this discrepancy can be explained by our HPLC results that either genetic or pharmacological reduction of DBH activity lowered NE abundance, but only *Dbh−/−* mice had supranormal DA levels. Because we and others have reported that nepicastat increases central DA concentrations in unstressed rodents (Devoto et al., 2015; Schroeder et al., 2010), we speculate that after exposure to a stressor, this increase in central DA is masked by increases in stress-induced DA synthesis, as we observed in this study (Abercrombie et al., 1989; Finlay et al., 1995; Sullivan and Gratton, 1998; Valenti et al., 2011).

The PFC, which receives catecholaminergic input from the LC and VTA, is implicated in emotional reactivity and behavioral control (McKlveen et al., 2015; Shansky and Lipps, 2013). Excitatory glutamatergic neurons in the PFC express receptors for catecholamines and project back to the LC and VTA, allowing for reciprocal communication with catecholaminergic nuclei (Kenwood et al., 2022; McKlveen et al., 2015; Shansky and Lipps, 2013). Hyperactivity of the PFC and its output regions in the striatum and thalamus are thought to contribute to repetitive behavior in compulsive disorders and their associated animal models (Ahmari and Rauch, 2022; Campbell et al., 1999; Manning et al., 2019). We found that control mice exposed to cage change with novel odorant stress had increased tissue concentrations of NE and DA in the PFC but not in the pons compared to mice exposed to the cage change only condition, suggesting enhanced catecholamine synthesis in the PFC (Fig. 6A). Importantly, phosphorylation of TH at residue Ser31, which increases TH activity (Dunkley and Dickson, 2019; Ong et al., 2014; Ong et al., 2011), was elevated in the PrL subregion of the mPFC but not the LC 30 min following novel odorant exposure compared to odorless cage change (Hwa et al., 2019; Ong et al., 2014), likely accounting for the higher NE and DA levels selectively observed in the PFC (Fig. 6B).

**Fig. 6.**
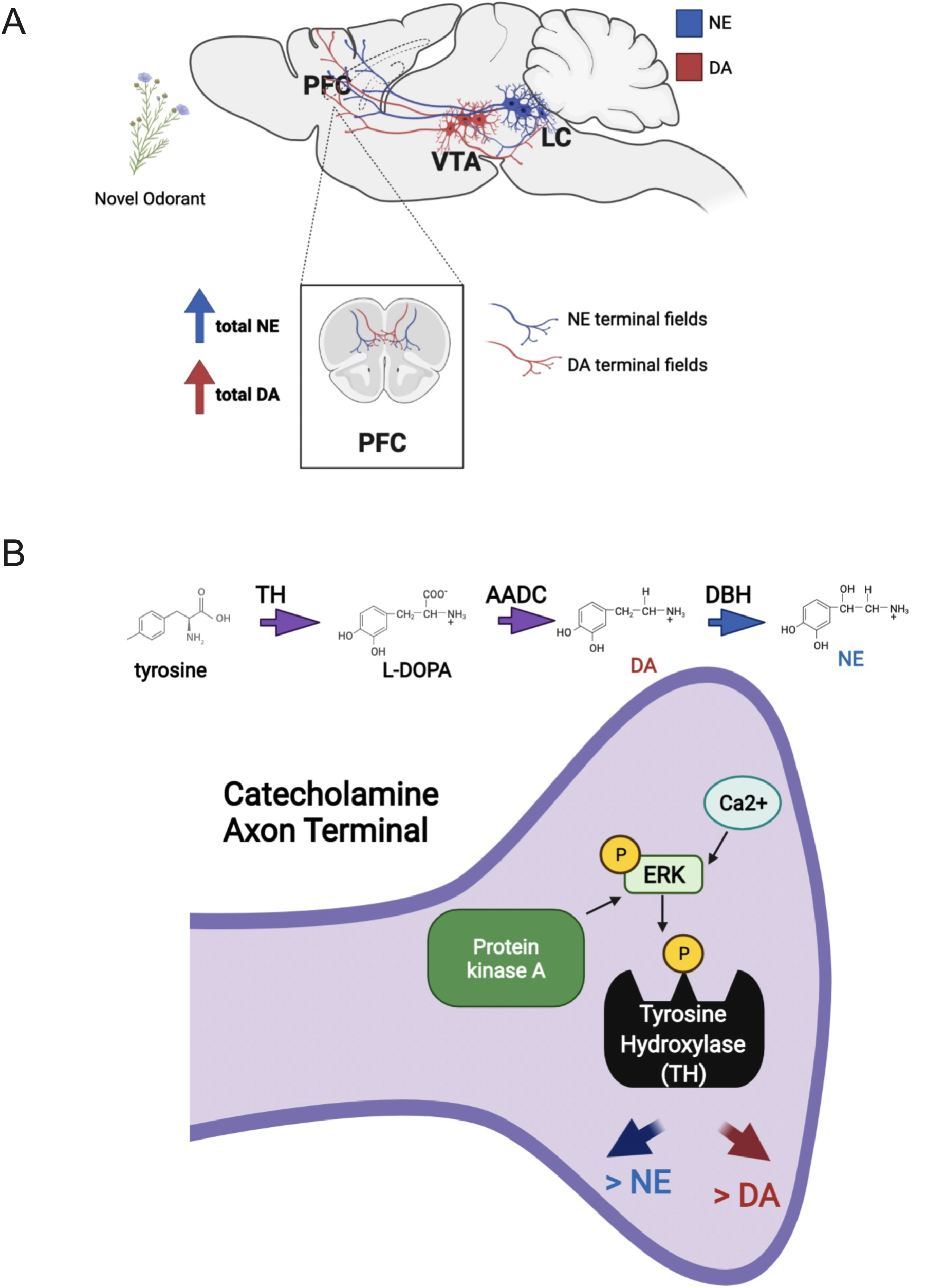
Summary of the effects of novel odorant stress on augmentation of catecholamine levels in the PFC. The PFC receives DA (red) and NE (blue) inputs from the VTA and the LC, respectively **(A)**. Both DA and NE neurons express tyrosine hydroxylase (TH), the rate-limiting enzyme in the synthesis of the catecholamine precursor L-DOPA; both cell types (indicated by purple coloring) also express the enzyme aromatic amino acid decarboxylase (AADC), which converts L-DOPA into DA. DA is converted to NE by the enzyme DBH in NE neurons, and DA neurons do not express DBH. Phosphorylation of TH Ser31 by activity-dependent kinases like ERK could result in rapid increases in the synthesis of both catecholamines in axon terminals within the PFC in response to stress **(B)**. *Created using Biorender.com*.

Catecholamine tone in the PFC and other forebrain regions involved in affective responses to stress fluctuate following exposure to various types of physical and psychological stressors (McKlveen et al., 2015; Shansky and Lipps, 2013), and NE is required for stress-induced anxiety-like and compulsive behavior in other models (Howard et al., 2008; Lustberg et al., 2020a; Lustberg et al., 2020b; Tillage et al., 2021). Thus, the complete lack of NE in the PFC may underlie the absence of typical digging behavior following cage change and novel odorant exposure in the *Dbh−/−* mice, although dysfunction of other stress-sensitive brain regions may also contribute (Schank et al., 2006; Weinshenker et al., 2002a). Central DA levels were supranormal in *Dbh−/−* mice but did not significantly differ between odorant conditions, raising the intriguing hypothesis that the “saturated” DA levels in the PFC of *Dbh−/−* mice could explain the abnormal grooming behavior regardless of odorant stress *in Dbh* mutants (Guild and Dunn, 1982; Taylor et al., 2010).

In our study, plasma CORT levels did not differ between odorless cage change and cage change + novel odorant conditions and was similarly unaffected by genetic or pharmacological disruption of DBH. As such, plasma CORT levels did not mirror the dramatic behavioral and neurochemical differences observed following novel odorant exposure and changes in DBH status. Changes in plasma CORT levels may be subject to ceiling effects that limit the sensitivity of this measure as an index of psychological stress (MacDougall-Shackleton et al., 2019), and elevations in plasma CORT are unlikely to be the causal mechanism driving behavioral differences between genotype and treatment groups. Critically, these findings indicate that stressor-evoked increases in circulating CORT occur independently of NE in this paradigm, which is consistent with previous findings from our lab measuring stress-induced anxiety-like behavior in *Dbh−/−* mice using other stressors like foot-shock (Tillage et al., 2021). In the same study, we showed that there are no differences in plasma CORT between *Dbh−/−* and *Dbh+/−* mice under basal conditions, nor differences in evoked CORT or freezing behavior during foot-shock.

We conclude that simple changes in behavior and PFC catecholamine content may be more sensitive measures of moderate psychological stress than plasma CORT levels in mice (Damanhuri et al., 2012; Iimori et al., 1982; Ong et al., 2011). In addition, using different plant-derived odorants in the same cohort of animals to maintain olfactory novelty, we have established a repeatable model of novel odorant stress that elicits defensive burying in NE-competent mice. Novel odorant stress may be considered alongside other stressful paradigms in which defensive burying is used as a behavioral readout, including predator odor exposure, marble burying, and the shock probe test (De Boer and Koolhaas, 2003; Hwa et al., 2019; Tillage et al., 2020b).

Our results have important implications for neuropsychiatric disorders. Digging and grooming are both repetitive behaviors which are measured in animal models of compulsive disorders like TS and OCD as analogs of compulsive behavior (Pittenger, 2015; Witkin, 2008; Wolmarans et al., 2016). Indeed, compulsive symptoms in patients with these conditions are exacerbated by stress, or perhaps even precipitated by it (Billnitzer and Jankovic, 2020; Chappell et al., 1994; Conelea and Woods, 2008). PFC function is critical for stress reactivity and behavioral control, and its dysregulation is thought to contribute to the pathophysiology of compulsive disorders (Ahmari and Rauch, 2022; Billnitzer and Jankovic, 2020; Shansky and Lipps, 2013).

Integrating our findings with the literature, we propose that excessive NE signaling within the PFC following psychological stress amplifies compulsivity in people with TS and OCD, and drugs which block NE transmission signaling may be useful for stress-related symptom exacerbation (Billnitzer and Jankovic, 2020; Nordstrom et al., 2015; Okazaki et al., 2019). Interestingly, administration of the α_1_AR antagonist prazosin or the inhibitory α_2_-autoreceptor agonist clonidine reduces tic behavior in the D1CT model of TS, which exhibit neuronal hyperactivity in the PFC that dysregulates the cortico-striato-thalamico-cortical (CSTC) loop (Godar et al., 2016; Nordstrom et al., 2015). The *Sapap3−/−* model of OCD, which also displays excessive grooming behavior and hyperactivity of the PFC, is associated with abnormalities in DA receptor expression in the CTSC loop (Ahmari et al., 2013; Manning et al., 2019; Manning et al., 2021). Plausibly, persistent excitation of PFC neurons by stress-induced catecholamine release may be a causal mechanism for increased repetitive behavior in patients with compulsive disorders and in animal models featuring excessive grooming as part of their phenotypes (Kasar and Yurteri, 2020; Leckman et al., 1995; McGrath et al., 1999; Zhang et al., 2013).

## Acknowledgments

We thank Synosia Therapeutics for providing the nepicastat and C. Brait for fruitful discussions regarding plant toxins and the adaptive value of innate aversion to floral odors in rodents. This work was supported by the National Institutes of Health (AG061175 and NS102306 to DW; NS007480-20 to DL; ES12870 to AFI).

## Notes

### Competing Interest Statement

DW is co-inventor on a patent concerning the use of selective dopamine beta-hydroxylase inhibitors for the treatment of cocaine dependence (US-2010-0105748-A1; Methods and Compositions for Treatment of Drug Addiction). The other authors declare no conflicts of interest.

### Summary of Updates

fixed changes with stars in graphs and with figure 5a label

